# BEACON: predicting side effects and therapeutics outcomes to drugs by Bridging knowlEdge grAph with CONtextual language model

**DOI:** 10.64898/2026.01.29.702277

**Authors:** Chengqi Xu, Jiaqi Xu, Krishna C. Bulusu, Heng Pan, Olivier Elemento

## Abstract

Biomedical knowledge graphs encode millions of relationships between drugs, proteins, pathways, and diseases, yet translating this structured knowledge into accurate predictions remains challenging. Existing deep learning approaches, including graph neural networks and knowledge graph embeddings, assign fixed representations to entities regardless of biological context, limiting their ability to capture how the same gene or pathway functions differently across scenarios. These methods also lack interpretability and often fail when applied to novel drugs outside their training distribution.

Here we present BEACON (Bridging knowlEdge grAph with CONtextual language model), a framework that transforms knowledge graphs into contextual sentence representations processable by language models. BEACON converts biomedical entities into tokens and relationships into syntactic dependencies, creating “sentence trees” that preserve graph structure while enabling contextual processing. A visibility matrix ensures that attention patterns respect the underlying knowledge graph topology, and a perturbation-based evaluation module identifies the specific genes, enzymes, and pathways driving each prediction.

We demonstrate BEACON’s versatility through two clinically important applications. For drug sensitivity prediction in cancer cell lines, BEACON achieves 0.941 AUROC and Spearman ρ = 0.919, outperforming existing methods (DrugCell, DeepCDR and DeepTTA). For drug-drug interaction (DDI) prediction, BEACON achieves 0.964 AUROC on the TwoSIDES benchmark and 0.84 AUROC on temporally held-out FDA adverse event data (2013–2023), demonstrating robust generalization to newly approved drugs. Applying BEACON to the BTK inhibitor acalabrutinib revealed that predicted interactions are enriched for drugs metabolized by CYP3A enzymes (OR = 3.01, P = 4.3 × 10⁻⁴), a mechanism validated through network proximity analysis. BEACON provides a unified, interpretable approach to knowledge graph-enhanced biomedical prediction.

## Introduction

Knowledge graphs (KGs) have revolutionized our ability to represent complex biomedical relationships, encoding millions of connections between drugs, proteins, pathways, and diseases in resources such as Hetionet, DRKG, and iBKH^1–6^. KGs can take many forms, e.g., drug-centric graphs capturing targets and metabolizing enzymes, gene-centric graphs encoding regulatory relationships, or disease-centric graphs linking phenotypes to molecular mechanisms. This structured knowledge provides a foundation for diverse biomedical applications: predicting drug efficacy or adverse interactions, identifying disease-gene associations, inferring gene function, or supporting clinical diagnosis. The common thread is translating a KG’s structured knowledge into predictions by leveraging the local neighborhood of entities being assessed.

Several deep learning approaches have been developed to leverage KGs for biomedical prediction, including graph neural networks and knowledge graph embedding methods^7–13^. However, these approaches face fundamental limitations. Current methods assign each entity, whether a gene, pathway, or drug, with a fixed embedding that remains unchanged regardless of biological context, preventing models from capturing how the same enzyme functions differently across metabolic scenarios. Graph neural networks and related architectures are also computationally complex and relatively inefficient. Critically, these methods fail to provide biological insight into the molecular mechanisms underlying their predictions, which is essential for clinical adoption and drug discovery.

To address these limitations, we present BEACON (Bridging knowlEdge grAph with CONtextual language model), a unified framework that transforms heterogeneous biomedical knowledge into contextual sentence representations. BEACON constructs “sentence trees” from KGs, where entities become tokens and relationships become syntactic dependencies. A GPT natural language backbone pre-trained on PubMed abstracts processes these representations, with custom visibility matrix constraints ensuring that attention patterns respect the underlying graph structure. This approach enables contextual entity representations, allowing the model to capture how biological components function differently across contexts, while maintaining explainability through perturbation-based importance scoring. Notably, BEACON employs a unified architecture adaptable to different prediction tasks by incorporating task-specific KGs while maintaining the same core methodology.

We demonstrate BEACON’s versatility through two fundamental challenges in precision medicine. First, drug sensitivity prediction—addressing why the same therapeutic produces dramatically different effects across cancer cell lines, which is essential for personalized treatment selection. BEACON achieves 0.941 AUROC in classifying sensitive cell lines, with interpretable predictions tracing from individual genes through pathways to cellular processes. Second, drug-drug interaction (DDI) prediction—critical given that DDIs account for 2-5% of hospital admissions and pose particular concern in oncology where polypharmacy is prevalent ^14,15^. BEACON achieves 0.964 AUROC on the TwoSIDES and data shift simulation benchmarks and shows strong zero-shot generalization (0.84 AUROC) on newly curated FDA adverse event data. Both applications share a common biological foundation: drug sensitivity and interactions arise from shared targets, enzymes, and pathways encoded in KGs. We applied BEACON to analyze acalabrutinib interactions, revealing CYP3A-enriched mechanisms validated through network proximity analysis. BEACON’s evaluation module identifies the specific KG components—genes, enzymes, pathways—that drive each prediction, enabling interpretable insights for precision medicine.

## Results

### A unified framework for KG-based biomedical prediction

Biomedical knowledge graphs encode rich relational information, connecting drugs to targets, enzymes to pathways, and genes to diseases, that can inform diverse prediction tasks. However, translating this structured knowledge into accurate predictions remains challenging. Current deep learning approaches assign each entity a fixed embedding that remains unchanged regardless of biological context, preventing models from capturing how the same gene or pathway functions differently across scenarios. We therefore aimed to design a unified framework that captures contextual relationships between biomedical entities while maintaining explainability of model predictions.

To capture biomedical KGs in a context-aware and interpretable model, we developed BEACON (Bridging knowlEdge grAph with CONtextual language model) as a framework with three modules: the knowledge module, the prediction module, and the evaluation module. The core idea is to transform hierarchical biomedical knowledge, whether drug-centric or cell-centric, into “sentence tree” representations that can be processed by language models while preserving the underlying graph topology.

The knowledge module is a knowledge encoder that constructs sentence trees from biomedical KGs, where each entity becomes a token and each relationship becomes a syntactic dependency. The sentence tree construction algorithm can be applied to distinct biological representations, such as drug-centered KGs and pathway-organized cellular states for response prediction (**Fig. 1**). The prediction module uses a GPT backbone pre-trained on 500,000 PubMed abstracts, processing sentence trees through transformer attention mechanisms. A custom visibility matrix constrains attention patterns to respect the underlying graph structure, so that entity representations are influenced only by their knowledge graph neighbors while benefiting from contextual language model processing. Task-specific prediction heads generate binary outputs for DDIs or continuous values for drug sensitivity. In addition to building highly accurate predictive models, the evaluation module employs an explainable approach to identify relevant knowledge-enriched features underlying prediction patterns, most notably an entity signature enriched in drug-centered KGs or a pathway signature related to cellular response.

**Fig. 1:**
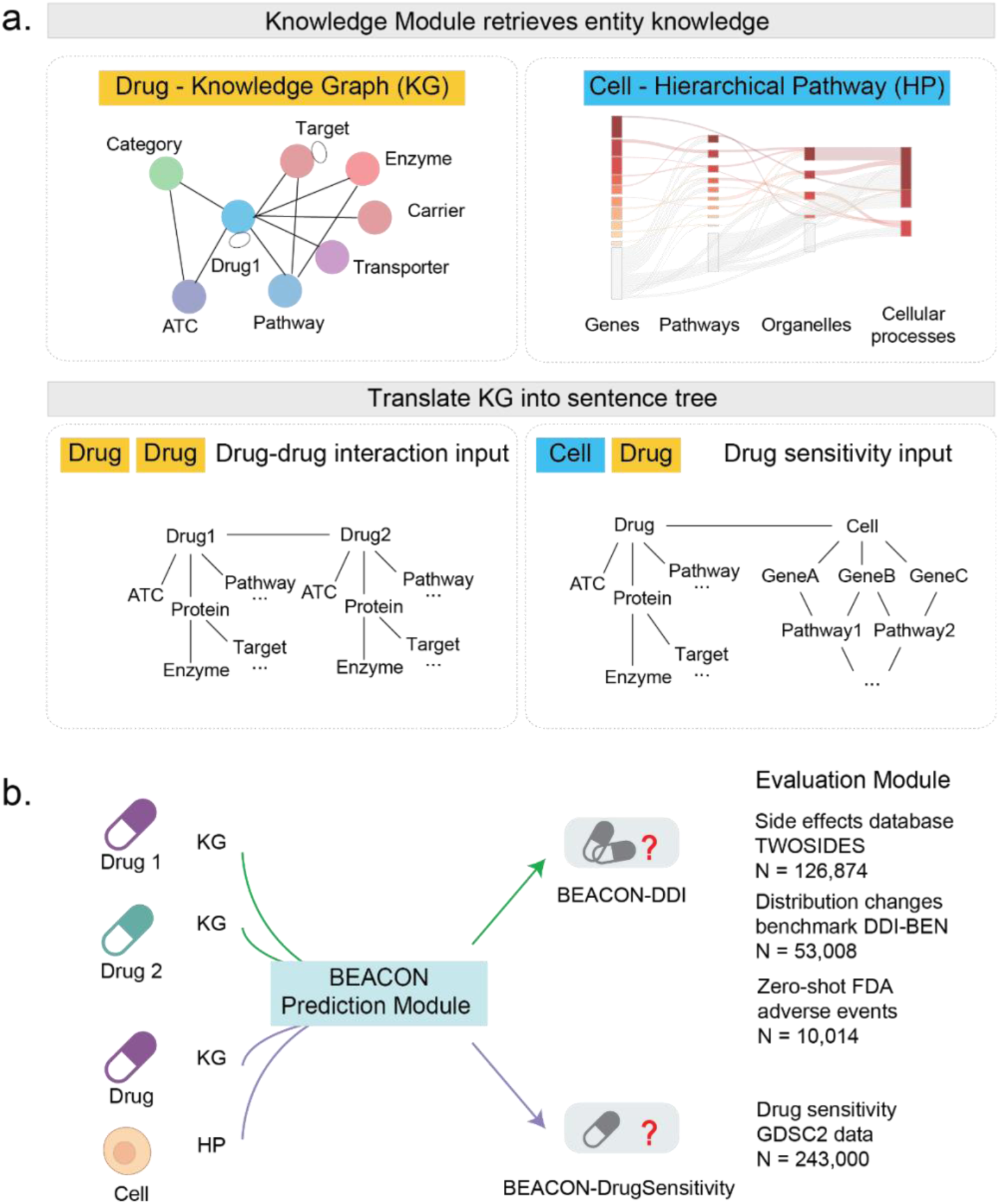
Schematic of the BEACON framework. **a.** Knowledge model by retrieving entity knowledge. BEACON integrates individual knowledge representations adapted to different entities. For example, a drug-centric KG encodes drug-associated entities, including targets, enzymes, carriers, transporters, pathways, and ATC classifications; and a cell-centric hierarchical pathway structure represents cellular states at multiple biological scales. Further, heterogeneous biomedical knowledge is transformed into sentence tree representations, where entities become tokens and relationships become syntactic dependencies. The hierarchical structure preserves relational semantics between connected entities. **b.** The prediction module takes natural language as inputs, which employs a GPT backbone with task-specific prediction heads to generate therapeutic outcome predictions. A transformer-based prediction module was implemented in BEACON for therapeutic outcome predictions (drug sensitivity or DDI).

### BEACON predicts drug sensitivity in cancer cell lines

A framework capable of predicting drug response variations across cell lines is valuable for patient stratification and personalized treatment selection (**Supplementary Fig. 1**). For drug sensitivity prediction, we constructed sentence trees from a cell-centric pathway hierarchy to generate “textual” representations for each cancer cell line. We collected pathway gene sets and hierarchical relationships from Reactome, where pathway relations P1 → P2 denote P1 as the child and P2 as the parent. This organization yielded 1,774 pathways spanning 4 hierarchical layers from small protein complexes to overarching cellular functions (**Methods**). Each cell line was represented by the top 15% most frequently differentially expressed genes, which were further organized into nested pathway sets representing cellular subsystems at different scales. The gene expression values were aggregated into pathway activity scores using GSVA, and the resulting pathway states were converted into sentence trees following the Reactome hierarchy. This representation enables multi-scale interpretation of drug sensitivity from individual gene contributions to pathway-level mechanisms.

To train the drug sensitivity prediction model, we harmonized data from the GDSC database, consisting of 809 cell lines and 265 drugs. BEACON was trained to associate each (cell line, drug) pair, represented by the cell’s pathway sentence tree and the drug’s KG sentence tree, with its corresponding drug sensitivity measured as IC50 values.

We then evaluated whether the resulting model (BEACON-DrugSensitivity) could predict drug sensitivity in cancer cell lines. We assessed prediction accuracy by comparing predicted and observed IC50 values. BEACON-DrugSensitivity achieved AUROC = 0.941 (95% CI: 0.932–0.950) for binary classification of sensitive cell lines (**Fig. 2a**). Performance exceeded DrugCell (AUROC = 0.917), which employs hierarchical pathway representations but relies on traditional neural network architectures rather than transformer-based contextual processing, suggesting that the contextual processing capabilities of language models provide additional predictive value beyond hierarchical knowledge organization alone. BEACON-DrugSensitivity also outperformed variant models variant models using biological features alone ( gene expression, mutation status) or drug structure alone (Morgan fingerprint), confirming that pathway-organized cellular states and drug knowledge graphs provide complementary predictive information (**Supplementary Table 1**).

**Fig. 2:**
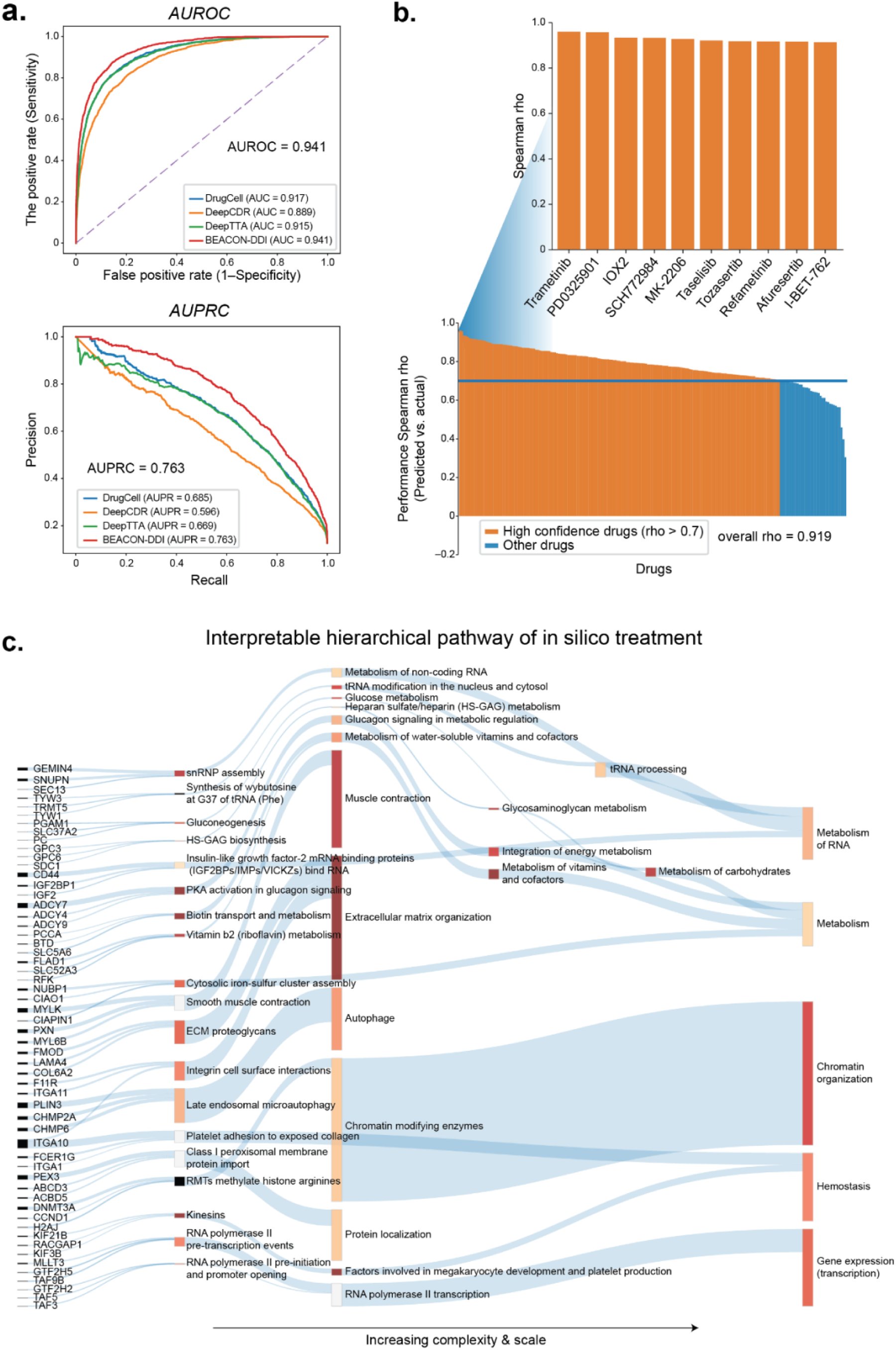
Overview of drug sensitivity prediction design. **a.** Comparison of predictive performances with state-of-the-art approaches DrugVNN, DeepCDR and DeepTTA. **b.** Drug-level prediction accuracy. Waterfall plot showing Spearman correlation (rho) between predicted and observed drug responses for each drug, ranked from highest to lowest. High-confidence drugs (rho > 0.7) are highlighted in red. **c.** Interpretable hierarchical pathway of in silico treatment. The multi-scale pathway hierarchy used to represent cell line states, spanning from individual genes (left) through large complexes and signaling pathways, to organelles and broad processes, to overarching cellular processes (right).

To assess drug-level prediction accuracy, we computed Spearman correlation for each compound individually. Of 265 drugs tested, 93 (35%) achieved high-confidence predictions (ρ > 0.7), with an overall correlation of 0.919 (**Fig. 2b**). Top-performing drugs included MEK inhibitors (trametinib, PD0325901, refametinib) and AKT inhibitors (MK-2206, afuresertib) and ERK (SCH772984) inhibitors.

To evaluate the relative importance of specific genes and pathways contributing to drug sensitivity predictions, we inspected each layer of the BEACON hierarchy and used a perturbation-based attribution method to obtain total importance scores (**Fig. 2c, Methods**). For each drug-cell pair, the model assigns an importance score (I, range 0–1) to each token, where higher values indicate greater influence on the prediction. For Trametinib, a MEK inhibitor, signaling genes such as *IGF2* and *ADCY7*, which mediate growth factor and cAMP crosstalk with the MAPK cascade, strongly contributed to predictions. In addition, highly ranked genes included *GTF2H5* and *TAF3*, which encode transcription machinery components downstream of MAPK signaling performance (**Supplementary Fig. 2a**). At the pathway level, importance converged on RNA polymerase II transcription and signal transduction processes across intermediate layers (**Supplementary Fig. 2b, c**), At the highest layer the top-ranked cellular processes were Gene expression (Transcription) and Metabolism (**Supplementary Fig. 2d**). Thus, the interpretable architecture of BEACON-DrugSensitivity can be interrogated to understand how the input information is transformed through layers and nodes, enabling further understanding of the state and importance of the involved biological entities.

### BEACON predicts DDIs with high accuracy

For DDI prediction, we leveraged a drug-centric KG to construct sentence trees encoding each drug’s biological neighborhood (**Supplementary Fig. 3a-d**). We then benchmarked the resulting model, BEACON-DDI against state-of-the-art methods and validated it through a case study of acalabrutinib interactions (**Supplementary Fig. 3e-g**). Here, we used a KG curated from the integrative Biomedical Knowledge Hub (iBKH). The KG encompasses 129,361 entities across multiple biological scales, from molecular targets and metabolizing enzymes to signaling pathways and therapeutic categories, along with 4,033,682 relationships (**Supplementary Table 2**). For each drug, the sentence tree encodes its associated targets, enzymes, carriers, transporters, and downstream pathway annotations, preserving the relational semantics between these entities, The drug’s KG neighborhood is into a hierarchical text structure. A detailed example of sentence tree construction is shown in **Fig. 3**.

**Fig. 3:**
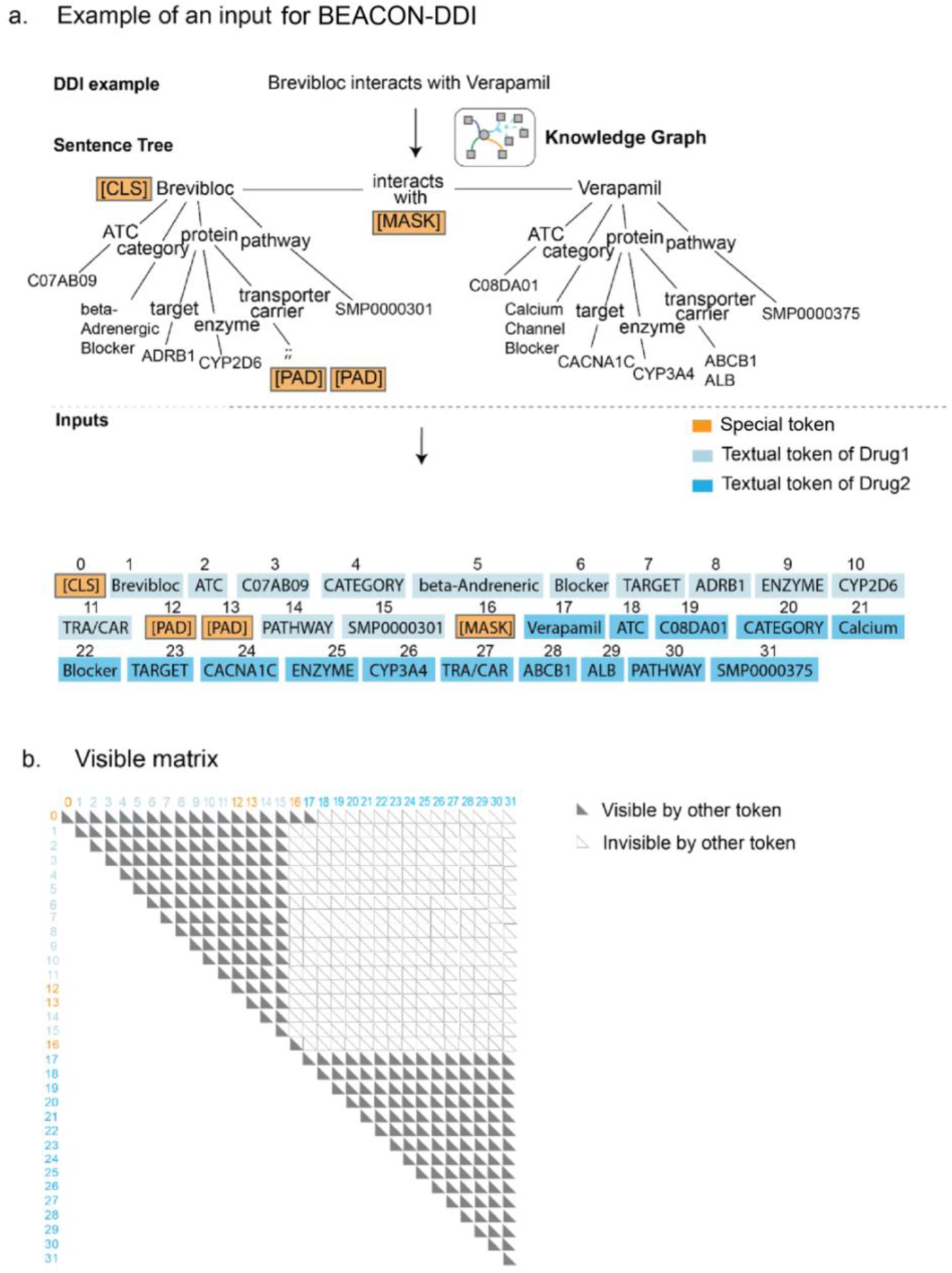
An example of a knowledge graph is enhanced to construct inputs for BEACON-DDI. **a.** The process of converting a DDI event, based on KG, into a sentence tree. Then the sentence tree is serialized into a natural text representation. This serialization process starts with a sentence containing the first drug’s ATC, category, target, enzyme, transporter/carrier (TRA/CAR), and pathway. It then describes the second drug. The special padded token PAD is added to the empty position of a sentence to ensure the sentence length are same if there is absent feature. The special masked token MASK is an indicator to the model that DDI label should be neglected during the process of training. The special token CLS is used for the classification of DDI label. **b.** The visible matrix limits the visible areas of each token by two rules: tokens belonging to two different drugs are masked, but tokens from the same drug are visible to each other.

To train the drug-drug interaction prediction model (Beacon-DDI), we used the expert-curated TwoSIDES dataset comprising 645 drugs with 63,437 interacting and 63,437 non-interacting DDI pairs^10^. BEACON-DDI was trained to associate each drug pair’s combined sentence trees with its corresponding interaction label. Importantly, both tasks share identical core components: the sentence tree construction algorithm, visibility matrix implementation, and GPT-based contextual processing, differing only in their input representations and prediction heads.

We implemented standard benchmarking using the TwoSIDES dataset. We randomly shuffled DDI pairs and set aside 20% (*n =* 25,376) as a holdout set. Here, we compared BEACON-DDI against five state-of-the-art machine learning methods, including structural similarity approaches (DeepDDI), modeling with heterogeneous biomedical entities (Decagon), and advanced GNN methods such as weighted graph convolutional networks (MR-GNN), heterogeneous attention networks (SSI-DDI), and nonlinear encoder-decoder layers (CASTER)^7,12,13,16,17^.

We evaluated model performance using three metrics: AUROC, the area under the precision-recall curve (AUPRC), and Precision@50. All five state-of-the-art methods achieved AUROC values above 0.80, with CASTER performing best at 0.920 AUROC. BEACON-DDI surpassed this performance with an AUROC of 0.964. For Precision@50, which assesses the accuracy of the top 50 ranked predictions for interacting DDIs, BEACON-DDI achieved 0.890, representing a 16% improvement over existing benchmarks (**Table 1**).

**Table 1.** Validation of BEACON and five state-of-the-art methods on the TwoSIDES dataset.

| Method | Representation | Architecture | AUROC | AUPRC | Precision @50 |
| --- | --- | --- | --- | --- | --- |
| BEACON<br>(Our model) | Knowledge Graph<br>Augmented Sentence<br>Tree | GPT framework | **0.964±0.002** | **0.943±0.001** | **0.890±0.002** |
| DeepDDI<sup>12</sup> | Structural Similarity | Fully Connected<br>Neural Networks | 0.826±0.002 | 0.812±0.001 | 0.734±0.001 |
| Decagon<sup>7</sup> | Heterogenous<br>Knowledge Graph | Encoder and Decoder<br>Layers | 0.863±0.002 | 0.809±0.002 | 0.803±0.001 |
| MR-GNN<sup>13</sup> | Molecular Graph | Weighted Graph<br>Convolutional<br>Networks and LSTM | 0.845±0.003 | 0.8029±0.003 | 0.728±0.001 |
| SSI-DDI<sup>16</sup> | Chemical Substructure | Graph Attention<br>Network | 0.858±0.001 | 0.782±0.001 | 0.827±0.001 |
| CASTER<sup>17</sup> | Chemical Substructure | Encoder and Decoder<br>Layers | 0.920±0.005 | 0.935±0.001 | 0.887±0.002 |

To understand BEACON-DDI’s predictive capabilities, we compared BEACON-DDI with two alternative architectures commonly used in sentence classification using identical inputs. First, we trained a conventional recurrent neural network (RNN) and a convolutional neural network (CNN) on the same sentence tree representations^8,18^. Both models showed inferior performance compared to BEACON-DDI (**Table 2**), confirming that the transformer architecture with visibility constraints captures more information from knowledge graph features than previous neural network approaches.

**Table 2.**
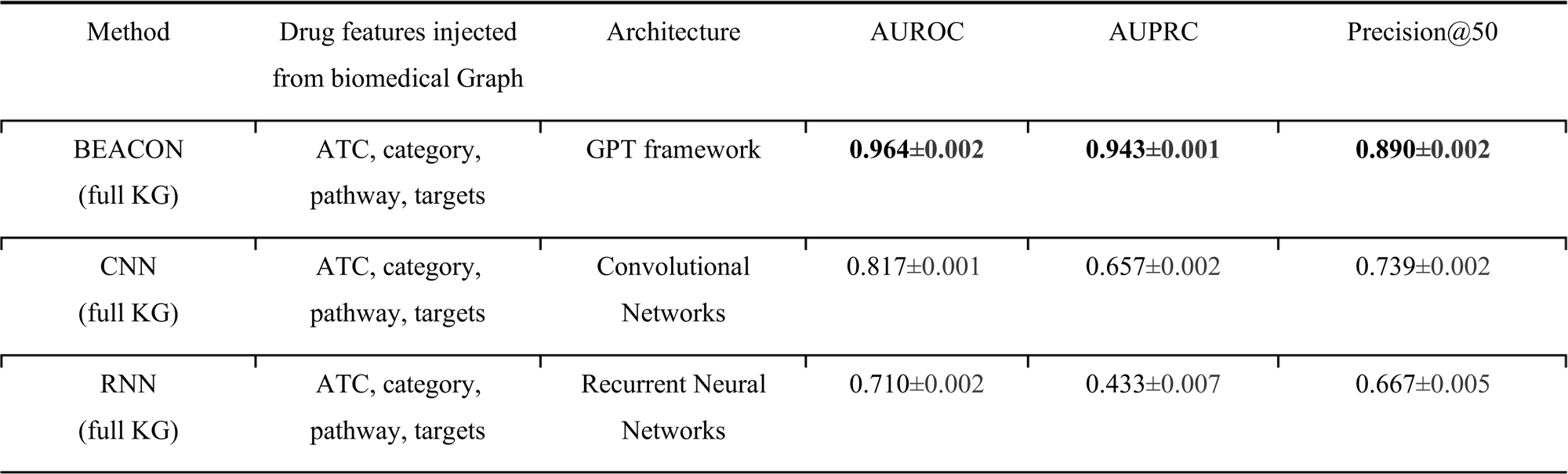
Prediction performance of BEACON versus the alternative deep learning models using the four input drug features injected from a biomedical KG.

We next evaluated how varying the components of KG background knowledge affects drug representations and DDI predictions. We conducted ablation studies using three KGs of different sizes, based on the number of biomedical entities included during training: Full KG (complete enriched dataset), BP KG (biological processes only), and PP KG (pharmacological properties only) (**Supplementary Note 1**). We found that BEACON-DDI with full KG features substantially outperformed both BP KG and PP KG variants (**Table 3**). Both BEACON-DDI and other deep learning models (RNN, CNN) showed optimal performance when using the full KG as input (**Table 3**), confirming that comprehensive KG representation of prior drug knowledge is critical for model performance.

**Table 3.** KG features are critical for model performance. On limited KG, BEACON still achieves robust performance.

| Method | Drug features injected<br>from biomedical Graph | Architecture | AUROC | AUPRC | Precision@50 |
| --- | --- | --- | --- | --- | --- |
| BEACON<br>(full KG) | ATC, category,<br>pathway, targets | GPT framework | **0.964±0.002** | **0.943±0.001** | **0.890±0.002** |
| BEACON<br>(ablated KG with<br>biological process<br>alone) | pathway, targets | GPT framework | 0.910±0.005 | 0.835±0.003 | 0.829±0.003 |
| BEACON<br>(ablated KG with<br>Pharmacological<br>properties alone) | ATC, category | GPT framework | 0.938±0.002 | 0.873±0.003 | 0.852±0.006 |

### Generalization to distribution shift scenarios

Predicting interactions for novel drugs remains challenging. When models rely on superficial patterns rather than deeper mechanistic understanding, they often fail to extend predictions to drugs lacking experimental data or complete KG information. This ‘shortcut learning’ phenomenon can lead to high performance on standard benchmarks but poor distribution shift prediction capability ^19^. Therefore, a model’s ability to infer missing links in KGs and make reliable predictions for drugs without prior interaction data is fundamentally important. However, for common DDI data split in most existing works ^7,20–22^, the phenomenon of distribution change is not considered, leading to unrealistic DDI evaluation results. Through the distribution change simulation framework, we can better simulate the distribution changes and conduct emerging DDI prediction evaluation in a scene closer to real-world scenarios. Therefore, we benchmarked BEACON-DDI against other predictive approaches on a new dataset under a recently proposed distribution changes framework^23^ (**Supplementary Fig. 4a**). BEACON-DDI maintained strong performance (AUROC = 0.91), substantially outperforming Decagon (0.78) and CASTER (0.82) (**Supplementary Fig. 4a**).

#### Validation on newly reported FDA data

The TwoSIDES training data was collected prior to 2013. To assess prediction of more recent DDIs, we curated an independent validation dataset from FDA Adverse Event Reporting System (FAERS) records^15^ spanning 2013Q1 to 2023Q2, yielding 9,480 confirmed DDIs after rigorous filtering (**Methods**). BEACON-DDI achieved an AUROC of 0.84 on this zero-shot prediction task, representing a 14% improvement over CASTER (AUROC = 0.74; **Supplementary Fig. 4b**). This result demonstrates that knowledge graph-based representations transfer effectively to DDIs arising from newly approved drugs.

### DDIs with acalabrutinib predicted by BEACON-DDI are enriched in drugs metabolized by the CYP3 family

After validating our model on an independent dataset, we applied BEACON-DDI to predict potential DDIs for acalabrutinib, a recently FDA-approved drug. The next-generation Bruton’s tyrosine kinase (BTK) inhibitor acalabrutinib was approved after our training dataset was released, making it an ideal candidate for evaluating zero-shot DDI prediction capabilities.

We performed a pairwise combinatorial assessment of 442 unique drugs from our independent dataset. Since acalabrutinib was absent from the initial training data, we updated the KG to incorporate comprehensive features for prediction purposes. Of the 442 drug combinations tested with acalabrutinib, 81 were predicted to have DDIs (aca-DDIs). We found that combinations involving CYP3A enzymes were enriched among DDI drugs with acalabrutinib (**Fig. 4a, Supplementary Fig. 5**). We further validated this finding through disproportionality analysis using the enzyme odds ratio (EOR) method, based on a 2x2 contingency table (**Supplementary Table 3, Methods**). The analysis demonstrated that drugs metabolized predominantly by CYP3A enzymes are 3.01 times more likely to cause DDI events when co-administered with acalabrutinib compared to drugs not metabolized by CYP3A (OR = 3.01, 95% CI: 1.57–5.78; P = 4.3 × 10^−4^; **Fig. 4b**).

**Fig. 4:**
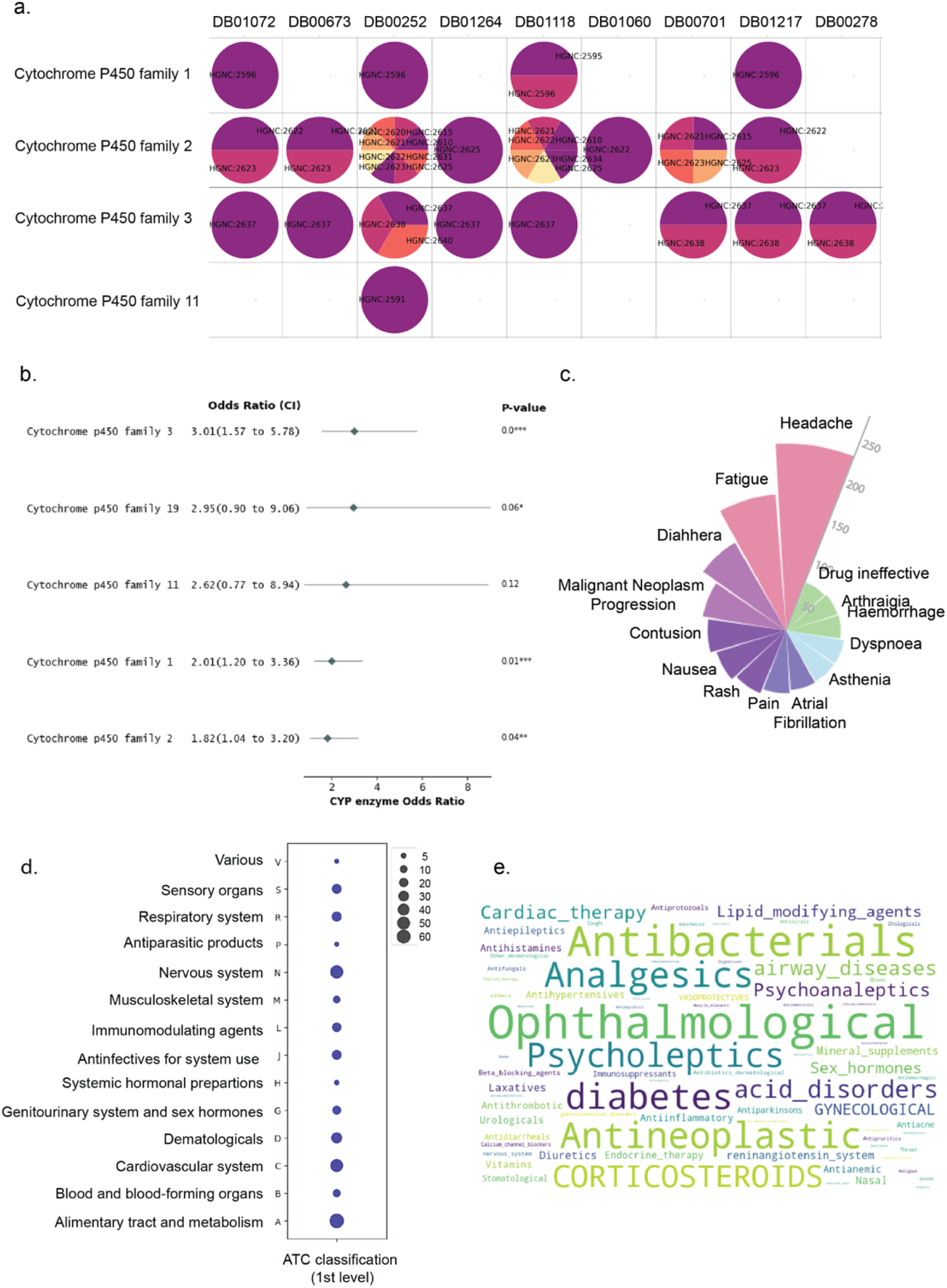
BEACON-DDI-predicted aca-DDIs are associated with the CYP3 enzyme family. **a.** Enzyme families of acalabrutinib predicted DDI drugs for disproportional analysis in **Supplementary Table 3**. The columns are drugs standardized with DrugBankID, the rows are the P450 enzyme family. Each circle indicates drug targets that belong to a certain family, presented by HGNC ID. If empty, this indicates that the drug has no targets metabolized by this enzyme family. **b.** Forest plot of the association of acalabrutinib DDI with the drug being metabolized by individual P450 enzymes. **c.** Common adverse event signals associated with predicted DDIs with acalabrutinib in the FAERS database**. d.** Distribution of ATC code of predicted DDIs belonging to the CYP3 enzyme family. **e.** Distribution of the therapeutic subgroup of predicted DDIs.

To characterize common phenotypes underlying aca-DDI drugs predicted to be substrates or inhibitors of CYP3A enzymes, we analyzed individual case safety reports (ICSRs) from FAERS where acalabrutinib was the primary suspect drug. Several common non-fatal side effects were observed across CYP3A-family DDIs when co-administered with acalabrutinib headache (*n* = 127 reports), fatigue (*n* = 98), diarrhea (*n* = 87), and malignant neoplasm progression (*n* = 64; **Fig. 4c**). We categorized these interacting drugs by their drug classes using first-level and second-level ATC codes. At the main pharmacological group level, nervous system, cardiovascular system, and alimentary and metabolism drugs were frequently reported to have DDIs with acalabrutinib (**Fig. 4d**). At the therapeutic subgroup level, antibacterial and chemotherapy drugs showed frequent DDIs with acalabrutinib (**Fig. 4e**). These findings highlight the diverse range of drug classes that may interact with acalabrutinib. The prominence of the alimentary tract and metabolism drugs aligns with the key role of hepatic CYP3A4, which metabolizes acalabrutinib and various other small-molecule drugs ^24^. Frequently co-prescribed medications including proton pump inhibitors and lipid modifiers often share this metabolic pathway, creating a chance of competition for enzyme binding and increasing the risk of reduced efficacy^25,26^. Similarly, the prevalence of nervous system and cardiovascular system reflects the clinical reality that patients with hematologic malignancies often require concurrent management of comorbidities (e.g., hypertension, pain) that require chronic management ^27^. Therefore, BEACON-DDI captures not only molecular interactions, but also real-world co-medication patterns, thereby bridging molecular mechanisms with potential clinical practice to proactively mitigate combination therapy risks.

#### Interpretable predictions through gene importance scoring

BEACON generates importance scores for each word in its analysis, revealing which elements most strongly influence its predictions. The evaluation module assesses these scores by introducing small controlled perturbations to input words and measuring how these changes affect DDI predictions. Words critical to the prediction show significant changes in DDI probability even with minimal perturbation.

To identify biologically meaningful patterns underlying DDI predictions, we analyzed importance scores across acalabrutinib DDIs using the perturbation-based approach (**Supplementary Fig. 6a, Methods**). Genes were clustered by importance values using unsupervised K-means clustering (*k* = 4), and pathway enrichment analysis was performed on each cluster using gene ontology (GO) terms (**Supplementary Fig. 6b**).

CYP3-associated aca-DDIs showed significant enrichment for genes involved in xenobiotic metabolism. The top-ranked pathways included cellular response to xenobiotic stimulus (FDR = 2.81 × 10^−4^) and xenobiotic metabolic process (FDR = 2.69 × 10^−3^; **Supplementary Fig. 6c**). BEACON-DDI captures mechanistically metabolic drug interactions at the enzyme level (CYP3A-mediated), as metabolic DDIs may require drug substitution for clinical management.

### Network analysis reveals mechanisms underlying predicted DDIs

In our previous study, PAIRWISE predicted 30 drugs as synergistic with acalabrutinib, which we validated through in vitro high-throughput screening ^28^. Among these, BEACON-DDI predicted 14 drugs (probabilities > 0.5) as having potential DDIs. To understand the mechanisms behind these predictions, we ranked genes by their importance scores and used network analysis to examine relationships between drug target networks and adverse reaction (ADR) networks. We obtained ADR gene sets from the ADReCS-Target database ^29^ and constructed drug target networks using BEACON-DDI’s highest-ranked genes. For each predicted DDI, we measured the network proximity between genes in the ADR subnetwork and the drug target subnetwork (**Fig. 5a-b**).

**Fig. 5:**
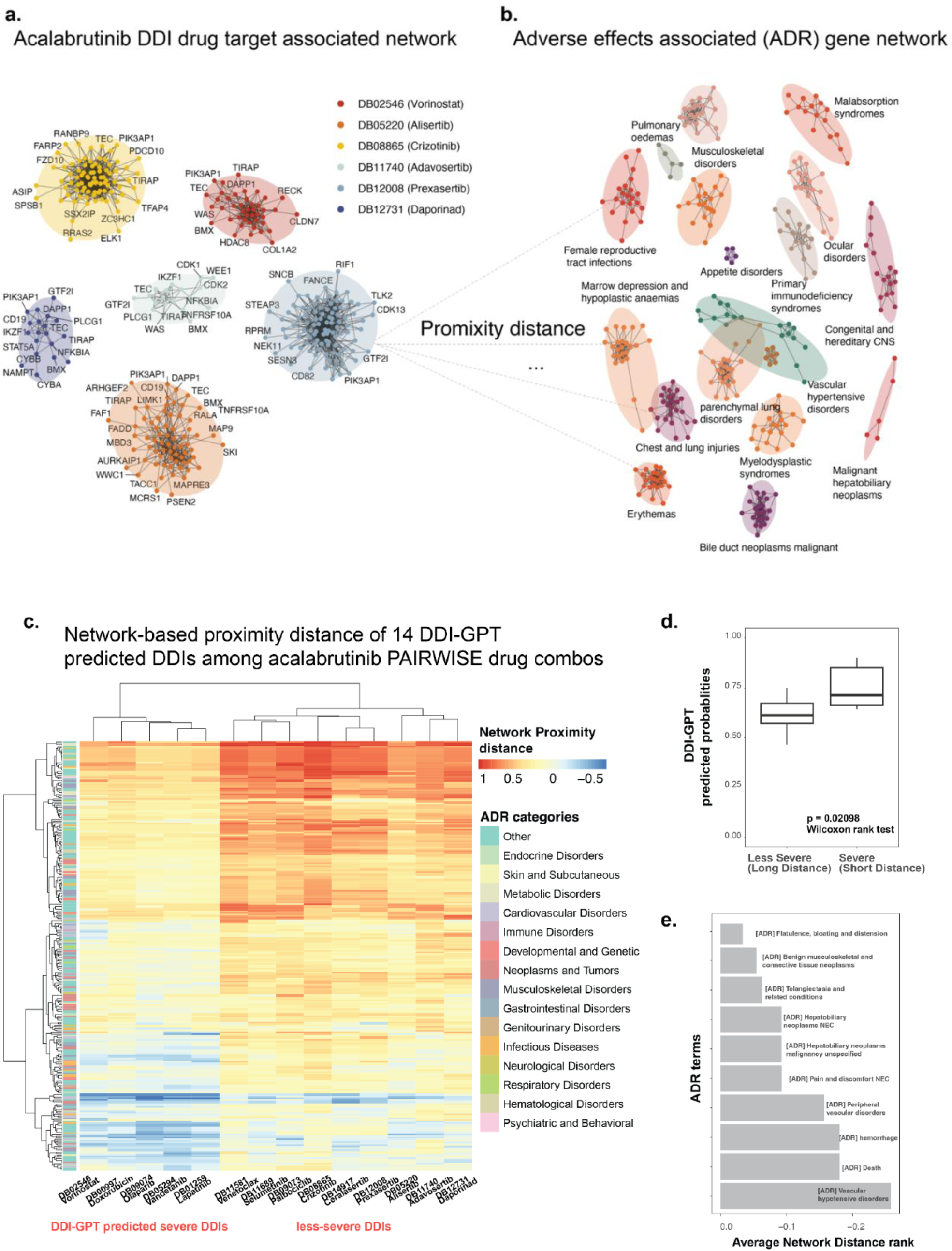
Network distance approach to identify DDI’s gene signatures. **a.** Each cluster in the graph on the left represents a single drug that is predicted as synergistic by the PAIRWISE model, and each node within a cluster represents important drug targets ranked by BEACON-DDI, with RWR on PPI. **b.** Each cluster in the graph on the right represents genes belonging to an ADR term. **c.** We calculated the network distance between individual drug target subnetworks and each ADR subnetwork. We plotted all combinations with their network distances, and unsupervised clustering revealed two distinct DDI profiles among 14 predicted DDIs, with the combination involving acalabrutinib. **d.** For two DDI profiles (severe or less-severe) created by paring 30 drug target subnetworks and 294 ADR subnetworks, we measured the correlation between the average distances of that profile and the BEACON-DDI predicted probability values for drug combinations of that profile. We found that combinations of drugs categorized by the distance approach tended to have higher BEACON-DDI predicted probability values in the severe group (*p* = 0.02098, Wilcoxon rank test). **e.** The top enriched ADR term in the set of genes contributing to DDI of drug combinations, including the acalabrutinib, sorted by their ranking of average network proximity distance across predicted DDIs.

Consistent with our hypothesis that drugs associated with CYP3A family would exhibit aca-DDIs, we found that strong CYP3A inhibitors (crizotinib, lapatinib, vorinostat) and moderate CYP3A inhibitors (olaparib, venetoclax, selumetinib) were highlighted by BEACON-DDI predictions, as noted by the Center for Drug Evaluation and Research (CDER), where these drugs require close monitoring when co-administered with acalabrutinib. Remarkably, we identified two distinct sets of drug combinations predicted as DDIs: combinations showing more severe effects when co-administered with acalabrutinib (characterized by shorter network distances) and combinations showing less severe effects (characterized by longer network distances) (**Fig. 5c**).

Measuring the relationship strength between average network proximity distance and BEACON-DDI predicted probability scores, we found that the less-severe group marked by shorter average network distance showed significantly increased DDI probability when co-administered with acalabrutinib (*p* = 0.002098, Wilcoxon rank test, **Fig. 5d**). One of the most enriched ADR terms for DDI is “vascular hypotensive disorders” (**Fig. 5e**). The genes associated with this ADR included BTK itself and downstream signaling molecules such as *PLCG2* and *MAPK1*, which play critical roles in vascular homeostasis and endothelial functions. Notably, drugs like lapatinib emerged due to their overlapping pathways and targets. For instance, lapatinib is a dual inhibitor of EGFR and HER2 tyrosine kinases and can influence the MAPK cascade, which is also modulated by acalabrutinib through BTK signaling. The convergence on the MAPK pathway suggests that co-administration could amplify effects on cell proliferation and vascular responses, potentially increasing hypotensive event risk. Additionally, the enrichment of “hepatobiliary neoplasms” suggests a possible link between acalabrutinib therapy and hepatic cellular processes (**Fig. 5e**). Genes like *CYP3A4* and *CYP3A5*, responsible for acalabrutinib metabolism in the liver, may contribute to hepatotoxicity or neoplastic transformations when their function is altered. Furthermore, the identification of “telangiectasia and related conditions” aligns with vascular anomalies that can arise from BTK inhibition. These findings highlight the utility of network analysis in linking drug target genes to enriched ADR terms.

## Discussion

In this paper, we have introduced BEACON, a unified framework that transforms heterogeneous biomedical knowledge graphs into contextual sentence representations for diverse prediction tasks. By bridging structured KG knowledge with the semantic capabilities of language models, BEACON addresses a fundamental limitation of existing approaches: the inability to capture how biological entities function differently across contexts. We demonstrate this framework’s versatility through two distinct applications — drug sensitivity prediction and drug-drug interaction prediction, both achieving strong performance while maintaining interpretability.

For drug sensitivity prediction, BEACON’s hierarchical pathway representations enable multi-scale interpretation, tracing predictions from individual genes through signaling pathways to cellular processes. This interpretable structure aligns with how biologists understand drug response: as an emergent property of pathway activities rather than isolated gene effects. The superior performance over models using biological features alone or drug structure alone confirms that integrating pathway-organized cellular states with drug knowledge graphs provides synergistic predictive power.

For DDI prediction, BEACON demonstrates superior zero-shot generalization on newly curated FDA data, a critical capability given that standard benchmarks may not reflect real-world performance on novel drugs. Our analysis of acalabrutinib interactions revealed CYP3A-enriched mechanisms, consistent with clinical pharmacokinetic studies. The gene importance scoring and network proximity analysis provide mechanistic insights that can guide clinical decision-making, identifying not just which drug combinations may interact, but also why. One limitation is that the enzyme family-DDI associations identified by our model cannot definitively establish causation; clinical relevance would require validation through in vivo trials.

The explainability of BEACON addresses a critical gap in biomedical AI. High accuracy alone is insufficient for clinical applications—providing clear explanations to patients, clinicians, and scientists is essential for establishing trust and clinical validity. BEACON’s evaluation module identifies the specific KG components driving each prediction, offering insights into complex drug-biology relationships at both pathway and interactome levels. This transparency distinguishes BEACON from “black box” approaches that achieve similar accuracy but offer no mechanistic insight.

Several directions could extend this work. First, incorporating personalized information such as patient transcriptome profiles and mutation status could enable truly individualized predictions. Second, as KGs are continuously updated and language models advance, BEACON’s architecture can incorporate richer representations from databases like PubChem and DrugBank. Third, real-world evidence from electronic health records could further validate and refine predictions. Finally, while we demonstrated BEACON on drug sensitivity and DDI prediction, the framework’s core methodology—transforming KG neighborhoods into contextual sentence trees—generalizes to other biomedical prediction tasks including gene function inference, disease-gene association, and clinical diagnosis support.

BEACON takes an important step toward interpretable, KG-enhanced biomedical prediction. By unifying diverse prediction tasks under a single architectural framework while maintaining explainability, BEACON enables both accurate therapeutic assessment and mechanistic insights for precision medicine.

## Methods

Our objective is to predict whether a combination of two drugs has adverse interactions with each other, particularly in a training scenario where limited samples are available. We first explain how we obtain the drug-related information from a heterogeneous knowledge graph (KG). The KG provides the language model with a structured approach to relate concepts. We then describe our proposed method that fine-tunes an language model-based model for DDI prediction.

### Curation of a Heterogeneous Knowledge Graph

We obtain heterogeneous graph representations of drugs by downloading a large-scale biomedical KG, iBKH, which covers all drugs in our benchmark TwoSIDES dataset and the external validation dataset, as well as 4 other types of biology entities, including drug targets, biological pathways, anatomical therapeutic chemicals, and drug categories, as well as the associated relations among them. Our KG is composed of entity-relation-entity triplets. Seven types of relations are used to connect the biomedical entities (**Supplementary Table 1**). We also constructed several KGs to represent different prior knowledge of drug entities, to test their effects on model performance (**Supplementary Note 1**).

### Integrating KGs into language model input

We used a two-step process to utilize a language model for biomedical graph data: knowledge query and knowledge injection. In the knowledge query step, all the biomedical entities related to a specific drug are selected to retrieve their corresponding triplets. In the knowledge injection step, the queried triplets are injected into sentence at their corresponding position, generating a sentence tree. The structure of the tree is illustrated in **Figure 2a**. In this study a sentence tree can have multiple branches.

#### Knowledge query

To retrieve all relevant information about a drug from its neighbors, we convert the structured graph into text for each instance of a drug. For example, given the biomedical entities (e.g., category, ATC, pathways, target, enzyme, carrier, transporter) and their relations with the input drug “*acalabrutinib*”, we extract a context-specific subgraph and transform it into a hierarchical sentence tree (Fig. 1a-b). The construction follows a depth-first traversal of the knowledge graph, with entities organized according to biological hierarchy: drug targets → metabolizing enzymes → carriers/transporters → associated pathways → therapeutic categories.

Each pathway is verbalized into a natural language description encoding its constituent genes and functional annotations. For example:

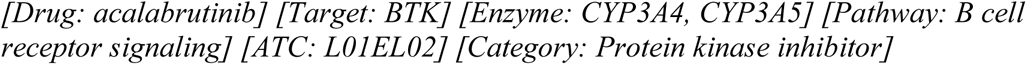

#### Knowledge injection

Direct concatenating a sequence of tokens within a knowledge graph triplet can potentially result in knowledge noise. In general, excessive incorporation of knowledge may divert the sentence from its intended meaning. Originally, K-BERT introduced soft-position and visible matrix to limit the influence ^30^. To overcome these challenges, inspired by K-BERT method, we propose a novel method to integrate KGs into DDI structure data via a sentence tree-based approach. Specifically, the DDI dataset is labeled at the single sentence level. Each sentence has a pair of drugs (*e1, e2*) with an annotated relation. We denote a sentence *s =* {*w_0_, w_1_, w_2_, …, w_n_*} as a sequence of tokens, where n is the length of this sentence. For an input DDI, we first transform it into the sentence “*Drug1* {*RELATION*} with *Drug2*”. The {*RELATION*} is set as a binary label {*interacting, non-interacting*} as ground truth for optimization during the training, and was not seen by BEACON in the input sentence, being replaced with a special MASK token. For an input sentence, the knowledge layer injects relevant triples from a KG into it, transforming the original sentence into a knowledge-rich sentence tree.

### BEACON model architecture

DDI prediction is a task to identify the association between drug pairs in an input sentence and assign the right classification to each pair. The task consists of three primary modules: the knowledge module, the prediction module, and the evaluation module.

#### Knowledge module

The knowledge module takes drug pairs as input, identifies their neighborhood in the KG, and transforms a drug pair into a knowledge-rich sentence tree. This sentence tree is subsequently translated into a token-level embedding representation. Since drug may have long or short dependencies with other drugs and biomedical entities, BEACON adopts self-attention mechanisms to capture the hidden long- and short-dependency information among different entities.

#### Prediction module

The classification module consists of a GPT-2 backbone optimized on a corpus of 500,000 PubMed abstracts and a linear layer for classification. GPT2 is developed with deep structures and pretrained textual data, which is the predecessor of GPT-3 and GPT-4. It achieved state-of-the-art results on several language modeling tasks, and with the smallest number of parameters (∼350M), which is applicable for fine-tuning with limited computation resources. In this study we applied the GPT-2 pretrained on 500,000 PubMed abstracts, referred to as BioGPT-2^11^. We built BEACON by fine-tuning BioGPT-2 with the TwoSIDES dataset, in order to adjust BioGPT-2 in the context of DDI prediction. To preserve the relational semantics of the knowledge graph during self-attention computation, we introduce a binary visibility matrix **M** ∈ {0,1}^n×n^ that constrains attention patterns:

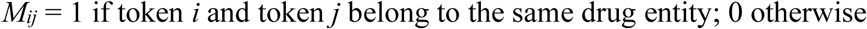

The masked attention score is computed as:

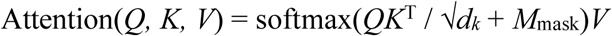

where *M*_mask_ = (1 − *M*). This design ensures that drug-specific features remain isolated during representation learning, preventing information leakage between drug entities in a pair.

To adapt the model for a binary-class classification task, we added a linear layer as a sequence classification on top of the pretrained BioGPT-2, which uses the special [CLS] token at the start of the model output to classify the DDI type. The same tokenizer used in BioGPT-2 was employed in BEACON. Cross-entropy loss was used to optimize the model during the fine-tuning process.

### Ranking important components in BEACON with evaluation module

To quantitatively determine the important input words in BEACON, we adopted a unified information-based measure described by Guan et al^31^. Briefly, a key task in explaining the black-box model is to associate latent representation with the interpretable input units (e.g., words) by measuring the contribution of the inputs. Existing methods largely rely on gradient-based methods; however, these can fail because they cannot explain how information flows through the network. In our study, we explain input importance through perturbation.

#### Perturbation-based importance

Following the unified information-theoretic framework, we assess pathway importance by measuring prediction sensitivity to controlled perturbations. For each input token *i*, we assign a perturbation weight σ_i_ ∈ [0,1] and generate noise vectors scaled by these weights:

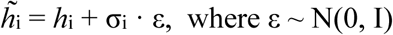

The importance score *I*_pert_(*i*) = 1 − σ_i_ reflects the noise tolerance of each token: tokens critical to predictions exhibit larger importance scores as they cannot tolerate perturbations without significant output changes. Pathway-level importance is obtained by averaging token-level scores within each pathway span.

### Statistical significance assessment

To determine whether pathway importance exceeds chance levels, we generate null distributions by randomly permuting gene-pathway memberships (*n* = 500 permutations) while preserving pathway sizes. We compute one-tailed *P* values and apply Benjamini-Hochberg correction for multiple hypothesis testing. Pathways with importance score ≥ 0.5 and FDR ≤ 0.1 are designated as core pathways for downstream biological validation.

### Experiments setting

#### Side-effects data

In this study, two real-world datasets were used to evaluate the BEACON performance. For the training, testing, and validation of the model, drug side effects are collected from the TWOSIDES dataset. To construct the external validation dataset for assessing the performance of our model, we adopted a new reference set of clinically relevant adverse DDI data^32^. We removed DDIs from this external validation set to ensure there were no overlapping data points seen during the training process. All curated datasets used in our study are provided in the supplementary data.

#### Curation of newly updated DDI data

We removed drug pairs that overlapped with the training data, yielding 9,480 unique DDIs associated with 442 distinct drugs. To create a negative control set, we randomly paired two drugs from the 442 unique drugs, ensuring the number of non-interacting drug pairs matched the number of interacting DDIs. To further verify that these randomly paired drugs were unlikely to interact, we conducted PubMed queries for each pair and excluded any that returned search results suggesting a known interaction. This rigorous process resulted in a final dataset of 13,586 drug pairs, including both interacting and non-interacting pairs, forming a comprehensive, updated dataset for independent validation of DDI predictions.

#### Metrics

Performance was evaluated using area under the receiver-operating characteristic (AUROC), area under the precision-recall curve (AUPRC) and average precision at 50 (AP@50). Higher values indicate better performance.

#### Evaluation strategies

We used the same data splitting procedure introduced by Zitnik et al^7^. For every experiment, we conduct five independent runs with different random splits of dataset. We then selected the best-performing model based on AUROC performance from the validation set. The selected model was then evaluated on the test set.

### Disproportionality analysis to determine CYP enzyme families associated with enriched DDIs predicted by BEACON

We propose using Enzyme Odds Ratio (EOR) to evaluate the possible association between a specific Cytochrome P450 (CYP) enzyme family and the DDIs predicted by BEACON This approach allows us to investigate whether a specific CYP enzyme family is enriched in DDIs more than would be expected by chance. There are 17 CYP enzyme families.

The EOR compares the odds of a specific CYP enzyme being associated with a particular DDI drug pair to all other enzymes, relative to the reporting odds for other drugs. We performed a disproportionality analysis to calculate the EOR. Before conducting this analysis, we first created a CYP enzyme contingency table (**Supplementary Table 3**) which served as the basis for the subsequent calculation of the EOR. The following formulas were used to calculate the EOR and its 95% confidence interval (CI):

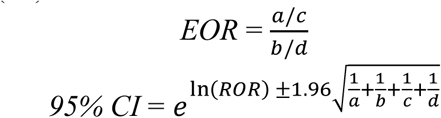

In these formulas, *a*, *b*, *c*, and *d* represent the counts from the contingency table. For the disproportionality analysis, we used the Chi-square test or Fisher’s exact test, and all statistical tests were two-tailed. All statistical analyses were conducted using Python SciPy v1.11.2 software.

### DDI-specific pathway analysis

To analyze the biological processes relevant to DDIs containing specific drugs in the dataset, we tested the top-ranked genes in the lists ordered by the importance values generated by the perturbation approach. We calculated pathway enrichments using GeneSet Variation Analysis (GSVA) R software package^33^ based on the importance value matrix for all predicted DDIs and non-DDIs containing the drug acalabrutinib. We used the set of pathways from Gene Ontology (GO) Biological Process terms for enrichment tests. The enrichments were calculated by a non-parametric estimator implemented in the ‘gsva()’ function. False discovery rate (FDR) correction was applied using the Benjamini-Hochberg procedure.

### Network distance measures

The adverse drug reaction (ADR) gene set was retrieved from ADReCS-Target (Huang et al., 2018), totaling 294 ADR phenotypes across 55,340 pairwise gene-ADR associations. The drug target gene set was retrieved from DrugBank database v 5.1.0^34^. A PPI network was obtained from STRING database v12.0^35^. We performed a Random Walk with Restart (RWR) algorithm to explore the global network structure of drug target on PPI interactome, using drug target as seed nodes to visit all nodes in PPI. The visiting nodes with higher-probabilities were used to complement and construct the drug target subnetwork. We employed network-based distance measures proposed by Cheng et al., (2019). In our context, given A, the set of ADR genes, T, the set of drug targets, and *d(A,T),* the closest distance measured by the average shortest path length between each node 𝑎 ∈ A and its nearest drug target 𝑡 ∈ T in PPI, the distance is defined as:

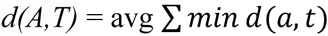

Here, *d(a,t)*represents the shortest path length between nodes 𝑎 and 𝑡.

### Development of standalone application

To support practical real-world application and assist domain experts in exploring DDIs predicted by BEACON, we have developed an interactive tool (**Supplementary Fig. 5).** This tool is a Python application coupled with the neo4j database that leverages our KG features at the backend. The frontend (i.e., the web application portal) was built using Streamlit v1.38.0, and NetworkX v.3.2.1 was used for graph visualization and data exploration. It is designed to support clinical decision-making and patient safety management. Users can search by entering a combination of two drugs, after which the backend framework automatically transforms these inputs into biologically meaningful sentences for BEACON prediction. Our pre-trained BEACON model then generates predictions for each DDI. The application also displays protein-protein and side-effect networks, which are visually represented as relational networks. Additionally, the application provides a table containing k-shortest path inference information from one to the other drug, as illustrated in the complex network, to facilitate analysis.

## Code availability

BEACON and the benchmarking platform can be downloaded from GitHub https://github.com/Mew233/ddigpt. Visualizations of BEACON web server (Supplementary Fig. 7) can be accessed on https://pyvisddi-24u28afk4upfhpvclyujvs.streamlit.app/

## Conflicts of interest

O.E. is supported by Janssen, J&J, AstraZeneca, Volastra, and Eli Lilly research grants. O.E. is scientific advisor and equity holder in Freenome, Owkin, Volastra Therapeutics and One Three Biotech and a paid scientific advisor to Champions Oncology. C.X., J.X., and H.P. declare no conflict of interest. KCB is an employee of AstraZeneca and holds shares in the company.

## Acknowledgement

The project was funded by AstraZeneca. O.E. is supported by UL1TR002384, UG3CA244697, R01CA194547, P01CA214274 grants from the National Institutes of Health and LLS SCOR grants 180078-02, 7021-20, 180078-01. H.P. was sponsored by Beijing Nova Program (20220484073). J.X. was supported by the Postdoctoral Fellowship Program (Grade B) of China Postdoctoral Science Foundation under Grant Number GZB20240044. The authors would also like to thank the wider collaborative teams from Weill Cornell Medicine and AstraZeneca for their support and valuable feedback right through the project.

## Supplementary material

### Supplementary Tables

**Supplementary Table 1.** KG features are critical for model performance. On limited KG, BEACON still achieves robust performance compared to CNN and RNN.

| Method | Drug features injected from biomedical Graph | Architecture | AUROC | AUPRC | Precision@50 |
| --- | --- | --- | --- | --- | --- |
| BEACON<br>(ablated KG with biological process alone) | pathway, targets | GPT framework | **0.910±0.005** | **0.835±0.003** | **0.829±0.003** |
| CNN<br>(ablated KG with biological process alone) | pathway, targets | Convolutional Networks | 0.762±0.002 | 0.601±0.005 | 0.702±0.004 |
| RNN<br>(ablated KG with biological process alone) | pathway, targets | Recurrent Neural Networks | 0.574±0.001 | 0.395±0.004 | 0.660±0.003 |
| BEACON<br>(ablated KG with Pharmacological properties alone) | ATC, category | GPT framework | **0.938±0.002** | **0.873±0.003** | **0.852±0.006** |
| CNN<br>(ablated KG with Pharmacological properties alone) | ATC, category | Convolutional Networks | 0.815±0.003 | 0.683±0.004 | 0.734±0.002 |
| RNN<br>(ablated KG with Pharmacological properties alone) | ATC, category | Recurrent Neural Networks | 0.577±0.004 | 0.401±0.002 | 0.670±0.003 |

**Supplementary Table 2.** Our curated KG is heterogenous, with 4 types of entities and 7 types of undirected edges. Tables below show a breakdown of entities by entity type and relations by entity type. **Supplementary Note 2** provides detailed information about the definition of entities.

| Entity pair | Relation types | Number of relations |
| --- | --- | --- |
| Drug-Gene | Drug-Target-Gene | 16,518 |
|  | Drug-Transporter-Gene | 3,066 |
|  | Drug-Enzyme-Gene | 5,241 |
|  | Drug-Carrier-Gene | 853 |
| Drug-Pathway | Drug-Association-Pathway | 3,231 |
| Drug-Category | Drug-Belongs to-Category | 599 |
| Drug-ATC | Drug-is_a-ATC | 599 |

**Supplementary Table 3.** Contingency table used for calculating EOR.

|  | Drugs predicted as DDIs with acalabrutinib | Drugs predicted as non-interacting with acalabrutinib | Total |
| --- | --- | --- | --- |
| CYP3 enzyme family | a | b | a+b |
| All other CYP enzyme families | c | d | c+d |
| Total | a+c | b+d | a+b+c+d |

a = The number of proteins belonging to CYP3 enzyme family across drugs which predicted to have DDIs with acalabrutinib b = The number of proteins belonging to CYP3 enzyme family across drugs which predicted to be non-interacting with acalabrutinib c = The number of proteins belonging to all other CYP enzyme families across drugs which predicted to have DDIs with acalabrutinib d = The number of proteins belonging to all other CYP enzyme families across drugs which predicted to be non-interacting with acalabrutinib

### Supplementary Note 1: Choice of graph to represent prior knowledge of the drug entity

We decided to proceed with the iBKH network to represent prior knowledge about the drug entities of interest when predicting co-prescribed drug combinations. The choice was made because iBKH network had the best coverage of the drug molecules we were interested in, and was the most general-purpose for application to future tasks. We studied the impact of varying the components of the curated KG that are used as prior knowledge.

#### Varying KG components

We first studied the impact of varying the components of the KG used as background knowledge for knowledge injection into the DDI sentence. We found that removing biological process-related entities, such as drug targets and pathways, resulted in the most significant decrease in performance This fits our assumption that drug targets are often critical for DDI prediction. Genes that share similar biological pathways should result in similar predicted DDI risk. While molecular function often overlaps with biological process annotation, biological process terms are more diverse and represent specific biological end states or outcomes. There was minimal impact on performance when pharmacological properties, such as ATC codes and categories, were removed.

#### Varying resolution of KG

Given a source drug, we query the KG by identifying its drug targets, transporters, and carrier proteins, with a maximum of *k = 2* of each node type. These *k* proteins are then connected to the source drug in the KG. By varying the *k* parameter, we can study the influence of the density of edges in the graph. We observed that the value of *k =2* resulted in the best performance of BEACON. When we reduced the value to *k =1*, thereby decreasing the density of the graph, we observed that the performance of BEACON decreased. Increasing the value to *k =3* or more had minimal impact on performance. We assume that genes that share similar biological pathways should result in similar predicted DDI risk.

### Supplementary Note 2

- *category*: This relation links drug nodes to MeSH (Medical Subject Headings) nodes.
- *ATC*: This relation links drug nodes to ATC (Anatomical Therapeutic Chemical) classification system code nodes.
- *pathway*: This relation links drug or protein nodes to pathway nodes.
- *target*: This relation links drug nodes to protein nodes.
- *enzyme*: This relation links drug nodes to protein nodes that catalyze chemical reactions involving the drug.
- carrier: This relation links drug nodes to protein nodes that are secreted proteins binding to drugs and carrying them to cell transporters.
- *transporter*: This relation links drug nodes to protein nodes representing membrane-bound proteins that shuttle ions into or out of cells.

**Supplementary Fig. 1:**
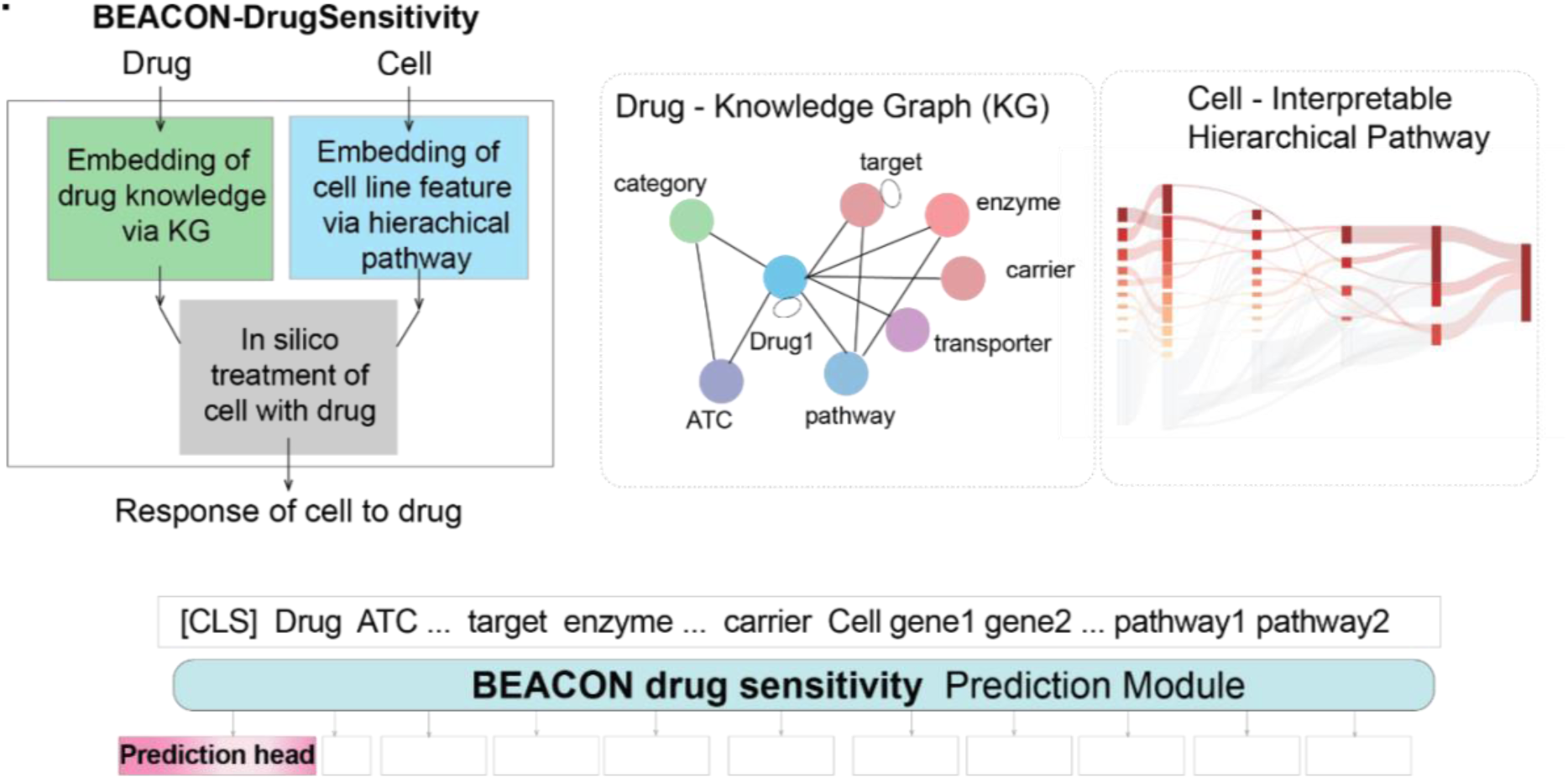
BEACON-DrugSensitivity predicts drug sensitivity by integrating drug embeddings from KGs with cell line embeddings from hierarchical pathways. Each drug is represented via its KG, while each cell line is represented through an interpretable hierarchical pathway structure derived from Reactome. These representations are concatenated as input tokens ([CLS] Drug ATC … target enzyme … carrier Cell gene1 gene2 … pathway1 pathway2) and processed by the BEACON drug sensitivity prediction module to generate response predictions.

**Supplementary Fig. 2:**
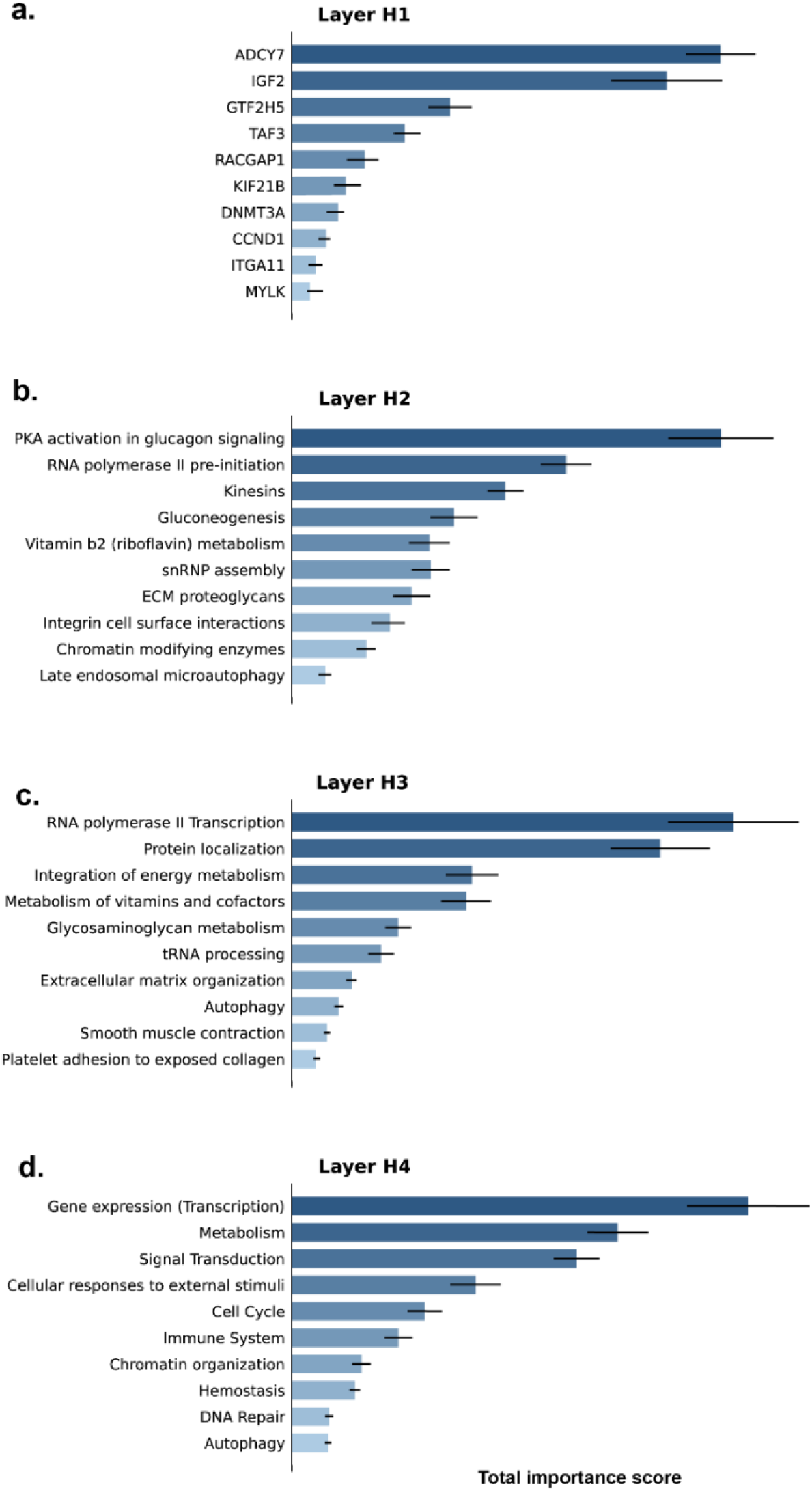
Relative ranking of nodes by total importance score for Trametinib sensitivity prediction. Horizontal bar charts showing the top 10–15 nodes in each layer ranked by total importance score, calculated as the summation of all sample-level importance scores over the testing set. Error bars represent the 95% confidence interval calculated using 1,000 bootstrap cycles.

**Supplementary Fig. 3:**
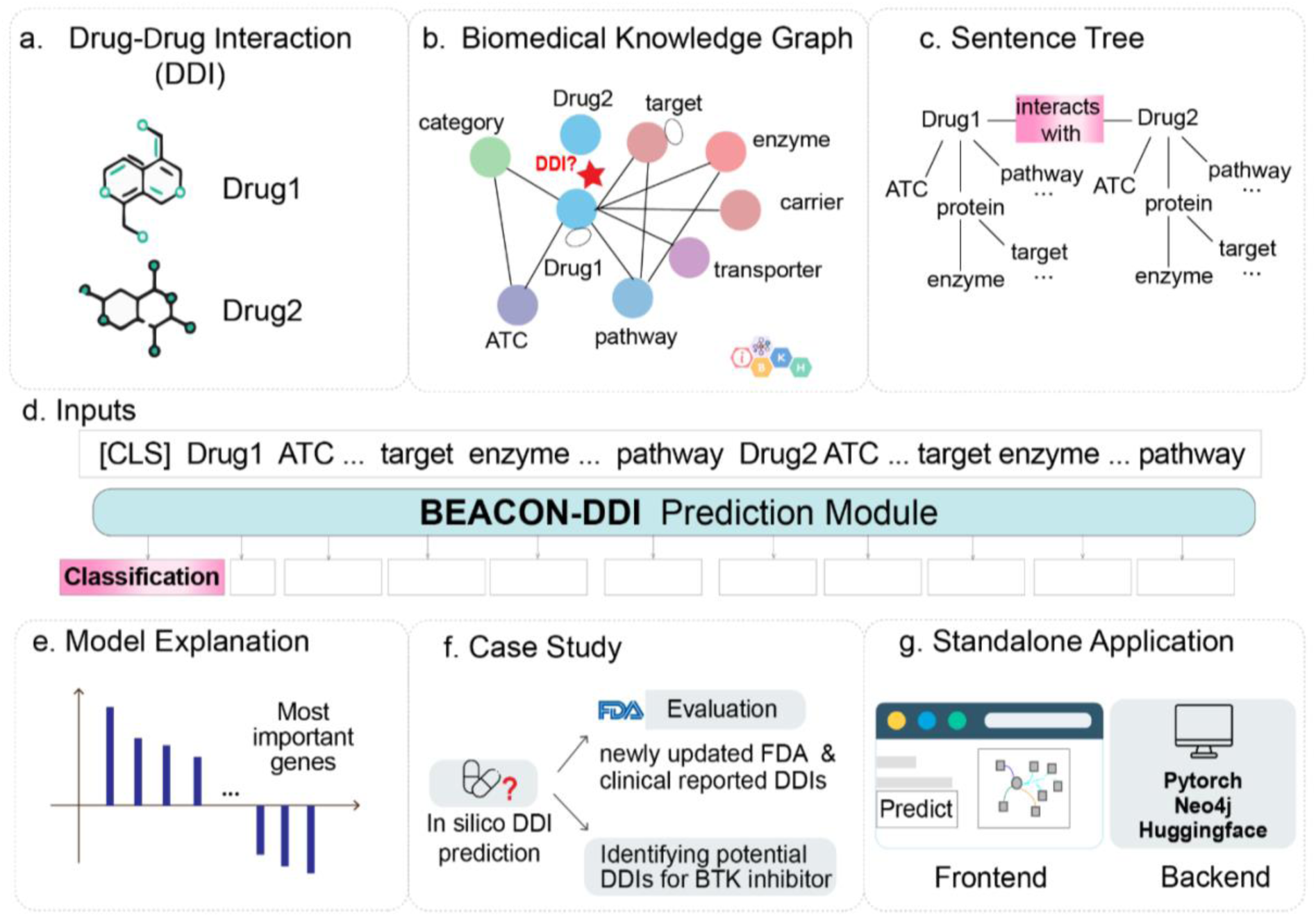
Overview of the BEACON framework on DDI prediction. **a.** Constructing drug-related multimodal representation through biomedical knowledge integration based on iBKH database. iBKH harmonizes data from 18 publicly available biomedical knowledge sources. We first collected diverse relation data which involves any drug entities in iBKH. Knowledge from diverse biomedical entities was integrated to build our curated drug-centered KG. **b.** We converted each DDI event into a sentence tree via knowledge injection from KG. **c.** The knowledge-enriched tree was translated into natural text and a task-specific prompt was created. The prompt was designed to generate binary class predictions of DDIs. **d.** A transformer-based prediction module was implemented in BEACON for novel DDI predictions. **e.** We explained the language model’ reasoning by ranking the most important genes nominated by BEACON-DDI. **f.** We validated our model on a new reference dataset which includes newly updated FDA and clinically reported DDIs. As a proof of concept, we analyzed *in silico* DDI predictions for the BTK inhibitor acalabrutinib. **g.** An interactive, dynamic web server was developed, which allows users to perform on-demand interface predictions using the BEACON-DDI framework.

**Supplementary Fig. 4:**
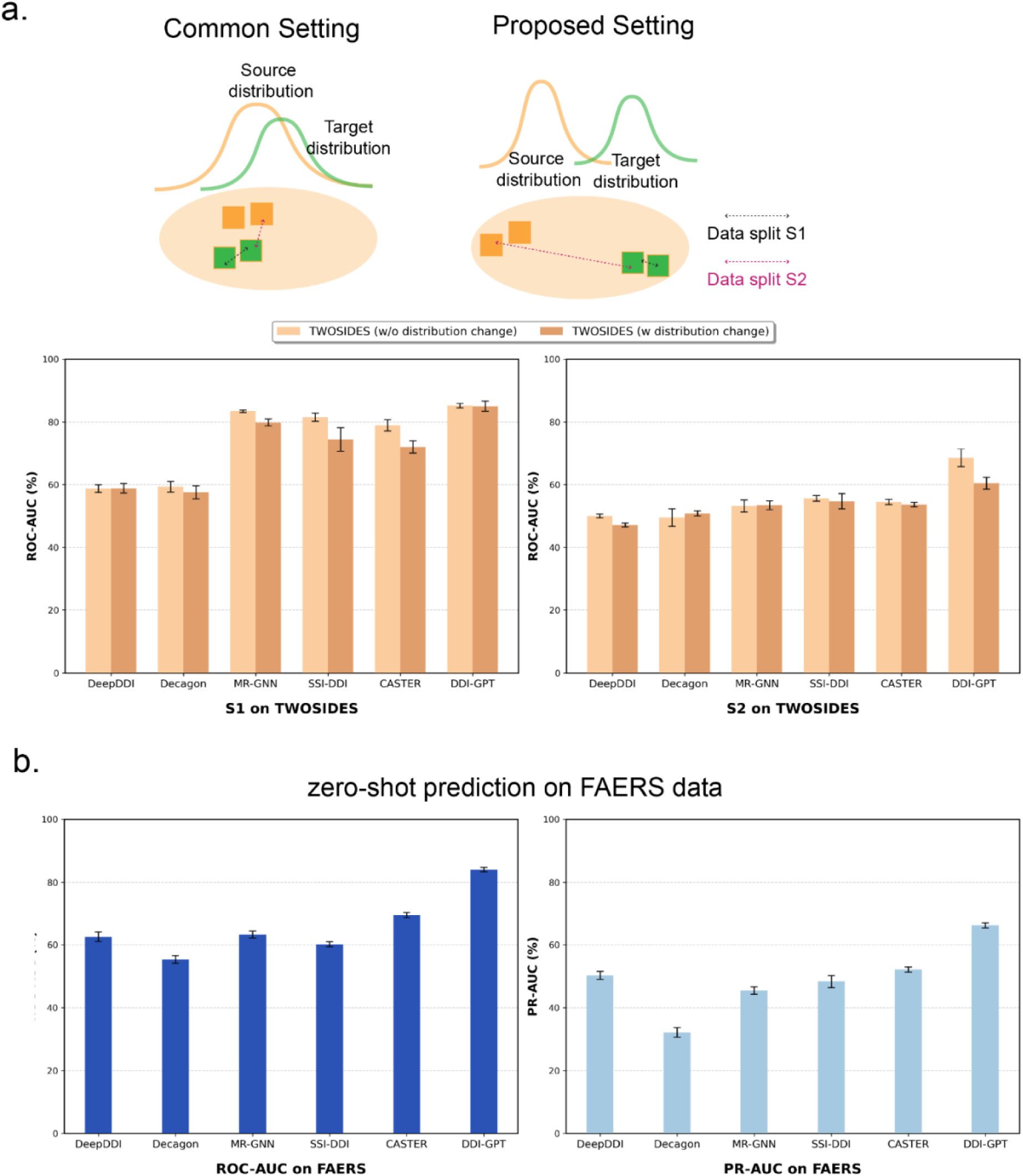
**a.** A data distribution framework ^23^ was employed to assess the model performance on generalization of new drug. We benchmarked DDI-GPT with other methods on the TWOSIDES dataset using two data splitting strategies where only one drug was unseen and two drugs were both unseen during the training. **b.** Predictive performances on an independent real-world. FAERS data, assessing the model’s zero-shot predictive capabilities.

**Supplementary Fig. 5:**
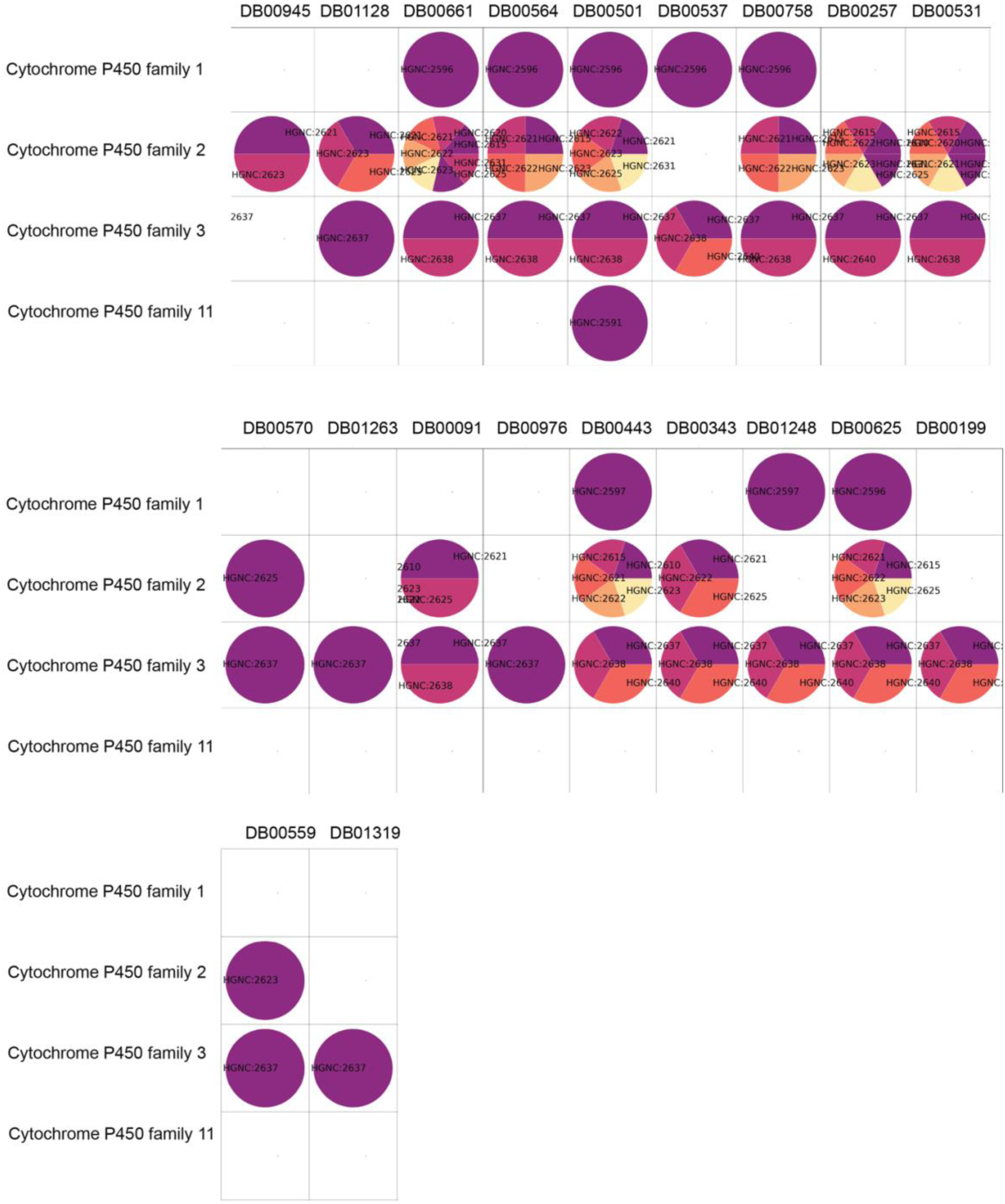
Enzyme families of acalabrutinib predicted DDI drug for disproportional analysis in **Supplementary Table 3**. The columns are drugs standardized with DrugBankID, and the rows are the CYP enzyme families. Each circle indicates a drug target that belongs to a certain family, presented by HGNC ID. If empty, this means that the drug is absent from targets metabolized by that enzyme family.

**Supplementary Fig. 6:**
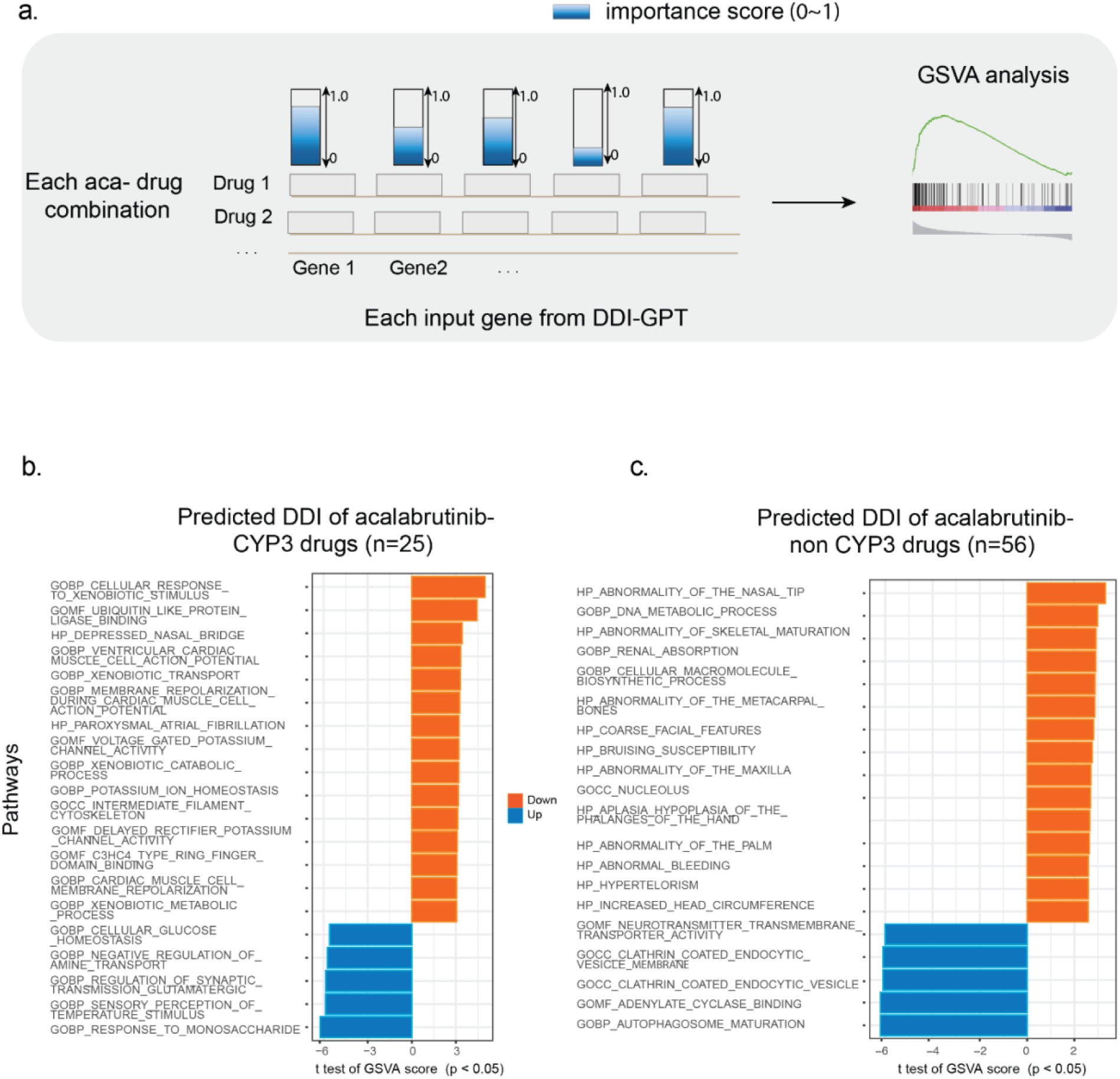
**a.** BEACON-DDI evaluation module uses importance scores to estimate the contribution of the word to the prediction, which can be computed efficiently by using perturbation-based approximation. The idea is to perturb the contribution of the word by adding a Gaussian noise and measure the magnitude of change in the prediction score. **b.** Genes are clustered based on their importance value by an unsupervised K-means clustering. Pathway analysis is performed on each cluster.

**Supplementary Fig. 7:**
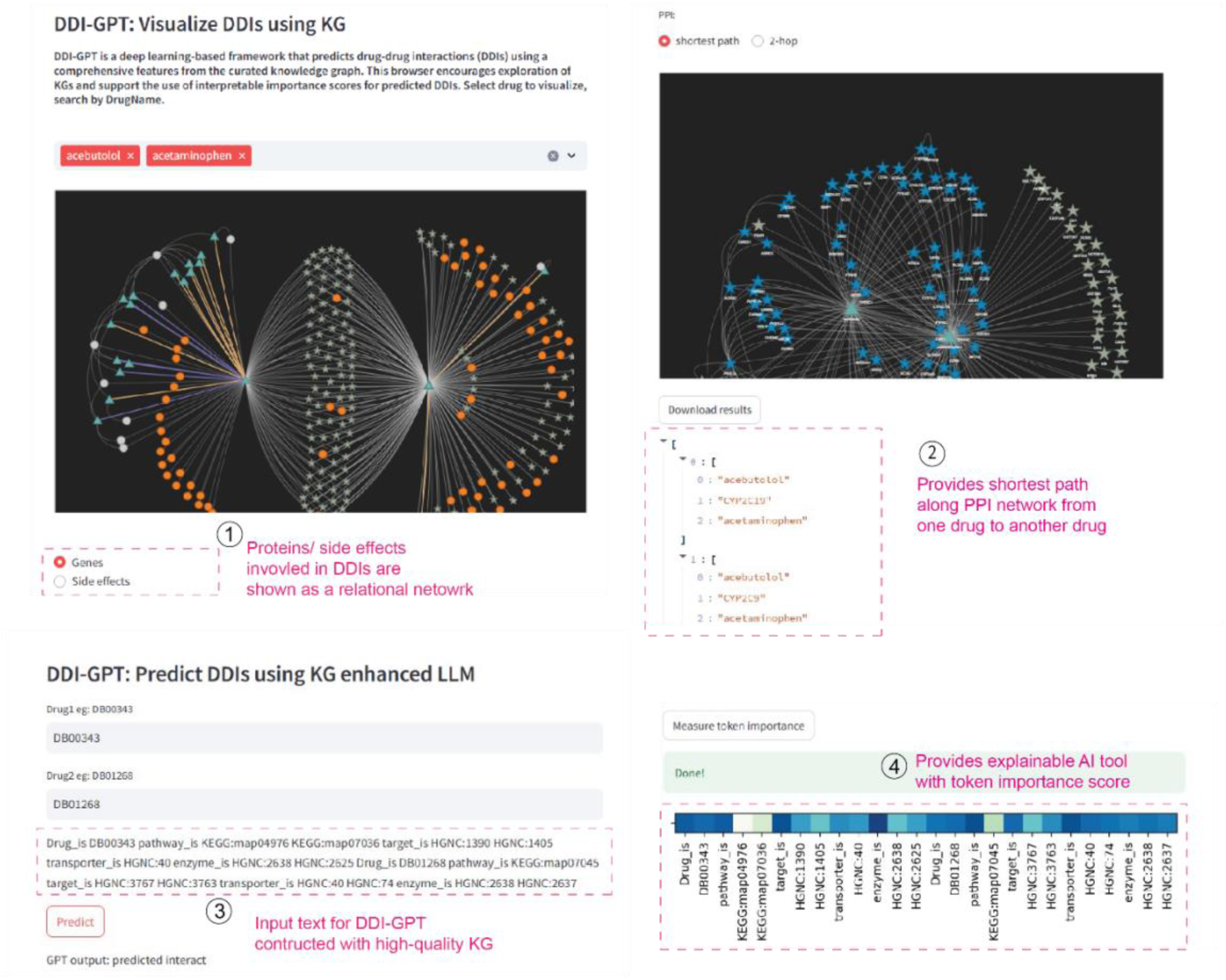
BEACON web server for knowledge-enhanced DDI exploration. The user submits a pair of drugs, referenced by DrugBank ID, and the input sentence is generated by incorporating drug-related biomedical entities from KG. The input is then set to a cluster running the prediction pipeline, where it is processed. Once the interface prediction is complete, the BEACON web server will be updated to visualize important words generated by the explanation module. The BEACON web server allows users to explore the KGs that were used by the BEACON framework, as well as various drug-protein-drug interaction networks and drug-side-effects-drug interaction networks. The user can view the shortest paths or two-hop paths between combinations of drugs.

## Notes

### Competing Interest Statement

The authors have declared no competing interest.

